# When null hypothesis significance testing is unsuitable for research: a reassessment

**DOI:** 10.1101/095570

**Authors:** Denes Szucs, John PA Ioannidis

## Abstract

Null hypothesis significance testing (NHST) has several shortcomings that are likely contributing factors behind the widely debated replication crisis of psychology, cognitive neuroscience and biomedical science in general. We review these shortcomings and suggest that, after about 60 years of negative experience, NHST should no longer be the default, dominant statistical practice of all biomedical and psychological research. Different inferential methods (NHST, likelihood estimation, Bayesian methods, false-discovery rate control) may be most suitable for different types of research questions. Whenever researchers use NHST they should justify its use, and publish pre-study power calculations and effect sizes, including negative findings. Studies should optimally be pre-registered and raw data published. The current statistics lite educational approach for students that has sustained the widespread, spurious use of NHST should be phased out. Instead, we should encourage either more in-depth statistical training of more researchers and/or more widespread involvement of professional statisticians in all research.

---

> *‘What used to be called judgment is now called prejudice and what used to be called prejudice is now called a null hypothesis. In the social sciences, particularly, it is dangerous nonsense (dressed up as the ‘scientific method’) and will cause much trouble before it is widely appreciated as such.’*
>
> — (Edwards, 1972; p. 180.)

> *‘…the mathematical rules of probability theory are not merely rules for calculating frequencies of random variables; they are also the unique consistent rules for conducting inference (ie. plausible reasoning)’*
>
> — (Jaynes, 2003; p.xxii)

## 1. The replication crisis and Null Hypothesis Significance Testing (NHST)

There is increasing discontent that many areas of psychological science, cognitive neuroscience, and biomedical research (Ioannidis 2005; Ioannidis et al. 2014) are in a crisis of producing too many false positive non-replicable results (Nosek et al. 2015). This wastes research funding, erodes credibility and slows down scientific progress. Since more than half a century many methodologists have claimed repeatedly that this crisis may at least in part be related to problems with Null Hypothesis Significance Testing (NHST) (Rozeboom 1960; Bakan 1966; Meehl 1978; Gigerezner 1998; Nickerson 2000). However, most scientists (and in particular psychologists, biomedical scientists, social scientists, cognitive scientists and neuroscientists) are still near exclusively educated in NHST, they tend to misunderstand and abuse NHST and the method is near fully dominant in scientific papers (Chavalarias, Wallach and Ioannidis, 2016). Here we provide an accessible critical reassessment of NHST and suggest that while it may have some legitimate uses NHST should be abandoned as the de factor *cornerstone* of research.

## 2. The origins of NHST as a weak heuristic and a decision rule

### 2.1 NHST as a weak heuristic based on the p value: Fisher

p values were widely popularized by Fisher (1925). In the context of the current NHST approach Fisher *only* relied on the concepts of the null hypothesis (H_0_) and the *exact p value* (hereafter p will refer to the p value and ‘pr’ to probability; see Appendix 1 for terms). He thought that experiments should aim to reject (or ‘nullify’; henceforth the name ‘null hypothesis’) H_0_ which assumes that the data demonstrates random variability according to some distribution around a certain value. Discrepancy from H_0_ is measured by a test statistic whose values can be paired with one or two-tailed p values which tell us how likely it is that we would have found our data *or* more extreme data if H_0_ was really correct. Formally we will refer to the p value as: pr(data or more extreme data | H_0_). It is important to realize that the p value represents the ‘extremeness’ of the data according to an imaginary data distribution assuming there is no bias in data sampling.

The late Fisher viewed the *exact* p value as a *heuristic piece of inductive evidence* which gives an indication of the plausibility of H_0_ together with other available evidence, like effect sizes (see Gigerenzer et al. 2004; Hubbard and Bayarri, 2003). Fisher recommended that H_0_ can usually be rejected if p ≤ 0.05 but in his system there is no mathematical justification for selecting a particular p value for the rejection of H_0_. Rather, this is up to the substantively informed judgment of the experimenter. Fisher thought that a hypothesis is demonstrable only when properly designed experiments *‘rarely fail’* to give us statistically significant results (Gigerenzer et al. 1989, p96; Goodman, 2008). Hence, a single significant result should not represent a ‘scientific fact’ but should merely draw attention to a phenomenon which seems worthy of further investigation including replication (Goodman 2008). In contrast to the above, until recently replication studies have been very rare in many scientific fields; lack of replication efforts has been a particular problem in the psychological sciences (Makel, Plucker and Hegarty, 2012), but this may hopefully change with the wide attention that replication has received (Nosek et al. 2015).

### 2.2 Neyman and Pearson: a decision mechanism optimized for the long-run

The concepts of the alternative hypothesis (H_1_), α, power, β, Type I and Type II errors were introduced by Neyman and Pearson (Neyman and Pearson, 1933; Neyman 1950) who set up a formal decision procedure motivated by industrial quality control problems (Gigerenzer et al. 1989). Their approach aimed to minimize the false negative (Type II) error rate to an acceptable level (β) and consequently to maximize power (1-β) *subject* to a bound (α) on false positive (Type I) errors (Hubbard and Bayarri, 2003). α can be set by the experimenter to an arbitrary value and Type-II error can be controlled by setting the sample size so that the required effect size can be detected (see Appendix 2 for illustration). In contrast to Fisher, this framework does not use the p value as a measure of evidence. We merely determine the critical value of the test statistic associated with α and reject H_0_ whenever the test statistic is larger than the critical value. The exact p value is irrelevant because the sole objective of the decision framework is long-run error minimization and only the critical threshold but not the exact p value plays any role in achieving this goal (Hubbard and Bayarri, 2003). Neyman and Pearson rejected the idea of inductive reasoning and offered *a reasoning-free inductive behavioural rule* to choose between two behaviours, accepting or rejecting H_0_, irrespective of the researcher’s belief about whether H_0_ and H_1_ are true or not (Neyman and Pearson, 1933).

Crucially, the Neyman-Pearson approach is designed to work efficiently (Neyman and Pearson, 1933) in the context of long-run repeated testing (exact replication). Hence, there is a major difference between the p value which is computed for a *single* data set and α, β, power, Type I and Type II error which are so called *‘frequentist’* concepts and they make sense in the context of *a long-run of many repeated experiments.* If we only run a single experiment all we can claim is that if we *had* run a long series of experiments we *would have had* 100α% false positives (Type I error) had H_0_ been true and 100β% false negatives (Type II error) had H_1_ been true *provided* we got the power calculations right. Note the conditionals.

In the Neyman-Pearson framework optimally setting α and β assures long-term decision-making efficiency in light of our costs and benefits by committing Type I and Type II errors. However, optimizing α and β is much easier in industrial quality control than in research where often there is no reason to expect a specific effect size associated with H_1_ (Gigerenzer et al. 1989). For example, if a factory has to produce screw heads with a diameter of 1±0.01 cm than we know that we have to be able to detect a deviation of 0.01 cm to produce acceptable quality output. In this setting we know exactly the smallest effect size we are interested in (0.01 cm) and we can also control the sample size very efficiently because we can easily take a sample of a large number of screws from a factory producing them by the million assuring ample power. On the one hand, failing to detect too large or too small screws (Type II error) will result in our customers cancelling their orders (or, in other industrial settings companies may deliver faulty cars or exploding laptops to customers exposing themselves to substantial litigation and compensation costs). On the other hand, throwing away false positives (Type I error), i.e. completely good batches of screws which we think are too small or too large, will also cost us a certain amount of money. Hence, we have a very clear scale (monetary value) to weigh the costs and benefits of both types of errors and we can settle on some rationally justified values of α and β so as to minimize our expenses and maximize our profit.

In contrast to such industrial settings, controlling the sample size and effect size and setting rational α and β levels is not that straightforward in most research settings where the effect sizes being pursued are largely unknown and deciding about the requested size of a good enough effect can be very subjective. For example, what is the smallest difference of interest between two participant groups in a measure of ‘fMRI activity’? Or, what is the smallest difference of interest between two groups of participants when we measure their IQ or reaction time? And, even if we have some expectations about the ‘true effect size’, can we test enough participants to ensure a small enough β? Further, what is the cost of falsely claiming that a vaccine causes autism thereby generating press coverage that grossly misleads the public (Godlee, 2011; Deer 2011)? What is the cost of running too many underpowered studies thereby wasting perhaps most research funding, boosting the number of false positive papers and complicating interpretation (Schmidt, 1992; Ioannidis 2005; Button et al. 2013)? More often than not researchers do not know the ‘true’ size of an effect they are interested in, so they cannot assure adequate sample size and it is also hard to estimate general costs and benefits of having particular α and β values. While some “rules of thumb” exist about what are small, modest, and large effects (e.g. Cohen, 1962; Cohen, 1988; Sedlmeier and Gigerenzer, 1989; Jaeschke et al. 1989), some large effects may not be actionable (e.g. a change in some biomarker that is a poor surrogate and thus bears little relationship to major, clinical outcomes), while some small effects may be important and may change our decision (e.g. most survival benefits with effective drugs are likely to be small, but still actionable).

Given the above ambiguity, researchers fall back to the default α=0.05 level with usually undefined power, these unjustified α and β levels completely discredit the originally intended *‘efficiency’* rationale of the creators of the Neyman-Pearson decision mechanism (Neyman and Pearson, 1933).

### 2.4. NHST in its current form

The current NHST merged the above concepts and is often applied stereotypically as a ‘mindless null ritual’ (Gigerenzer, 2004). Researchers set H_0_ nearly always ‘predicting’ zero effect but do not quantitatively define H_1_. Hence, pre-experimental power cannot be calculated for most tests which is a crucial omission in the Neyman-Pearson framework. Researchers compute the *exact* p value as Fisher did but also *mechanistically* reject H_0_ and accept the undefined H_1_ if p≤(α=0.05) without flexibility following the *behavioural decision rule* of Neyman and Pearson. As soon as p≤α, findings have the supposed right to become a scientific fact defying the exact replication demands of Fisher and the belief neutral approach of Neyman and Pearson. Researchers also interpret the *exact p* value and use it as a relative *measure of evidence* against H_0_, as Fisher did. A *‘highly significant’*result with a small p value is perceived as much stronger evidence than a weakly significant one. However, while Fisher was conscious of the weak nature of the evidence provided by the p value (Wasserstein and Lazar, 2016), generations of scientists encouraged by incorrect editorial interpretations (Bakan 1966) started to exclusively rely on the p value in their decisions even if this meant neglecting their substantive knowledge: scientific conclusions *merged* with reading the p value (Goodman, 1999).

## 3. Neglecting the full context of NHST leads to confusions about the p value

Most textbooks illustrate NHST by partial 2×2 tables (see **Table 1**) which fail *to* contextualize long-run conditional probabilities and fail to clearly distinguish between long-run probabilities and the p value which is computed for a single data set (Pollard and Richardson, 1987). This leads to major confusions about the meaning of the p value (see Box 1).

### BOX 1: Major confusions about the p value

1. Many practicing researchers and even some statisticians confuse the roles of the p value and α(Hubbard and Bayarri 2003). These researchers set α = 0.05 before they run an experiment but once they compute the p value they falsely assume that the p value will now represent the actual data-dependent Type I error probability somehow replacing the Neyman-Pearson α level while also interpreting it as the strength of evidence against H_0_ as used by Fisher (Goodman, 1993; 1999; Nickerson 2000). However, α is always fixed independently of what p value we find in an experiment whereas p values can be considered random variables, varying widely from experiment to experiment (Murdoch et al. 2008; Hung et al. 1997; Simonsohn et al. 2014a,b; Sterling 1959). Currently, the expression ‘significance level’ is used interchangeably for both the p value and α reflecting the confusion about them (Hubbard and Bayarri 2003).
2. Many practicing researchers falsely assume that if p = 0.01 then the probability of a false positive finding given the data (pr(H_0_|data) is 0.01. Conversely, they also assume that if p = 0.01 then the probability of a truly positive finding given the data (pr(H_1_|data) is 1 - p = 0.99. Yet, others confuse the p value with the ‘updated’ H_0_:H_1_ odds after a study was run, and/or with replication success (Bakan, 1966; Meehl, 1967; Pollard and Richardson, 1987; Cohen 1994; Hunter, 1997; Goodman, 1999; Oakes, 1986; Gliner et al. 2002; Wilkerson and Olson, 2010; Hoekstra et al. 2014; Castro-Sotos 2007; 2009). These *false* assumptions are not only *thoroughly wrong,* they also deeply *underestimate* the probability of false positive findings and highly *overestimate* the probability of truly positive findings and replication success. The network of confusions outlined here constitute what Goodman (1999) termed the 'p *value fallacy′* (see Goodman, 1999; Goodman 2008; Nickerson 2000 and Wagenmakers, 2007 for excellent reviews).

First, both H_0_ and H_1_ have some pre-study or ‘prior’ probabilities, pr(H_0_) and pr(H_1_). This means that before the study is run we may have some knowledge about the validity of H_0_ and H_1_. For example, we may know about a single published study claiming to demonstrate H_1_ by showing a difference between appropriate experimental conditions. However, in conferences we may have also heard about 9 highly powered but failed replication attempts very similar to the original study. In this case we may assume that the odds of H_0_:H_1_ are 9:1, that is, pr(H_1_) is 1/10. Of course, these pre-study odds are usually hard to judge unless we demand to see our colleagues’ ‘null results’ hidden in their drawers because of the practice of not publishing negative findings. Current scientific practices appreciate the single published ‘positive’ study more than the 9 unpublished negative ones perhaps because NHST logic only allows for rejecting H_0_ but does not allow for accepting it *and* because researchers *erroneously* often think that the single published positive study has a very small, acceptable error rate of providing false positive statistically significant results which equals α, or the p value. So, they often spuriously assume that the negative studies somehow lacked the sensitivity to show an effect while the single positive study is perceived as a well-executed sensitive experiment delivering a ‘conclusive’ verdict rather than being a ‘lucky’ false positive (Bakan, 1966).

NHST completely neglects the above mentioned pre-study information and exclusively deals with rows 2-4 of **Table 1**. NHST computes the one or two-tailed p value for a particular data set assuming that H_0_ is true. Additionally, NHST logic takes long-run error probabilities (α and β) into account conditional on H_0_ and H_1_ These long-run probabilities are represented in typical 2x2 NHST contingency tables but note that β is usually unknown in real studies.

As we have seen, NHST *never* computes the probability of H_0_ and H_1_ being true or false, all we have is a decision mechanism hoping for the best individual decision in view of long-run Type I and Type II error expectations. Nevertheless, following the repeated testing logic of the NHST framework, for many experiments we can denote the *long-run probability* of H_0_ being true given a statistically significant result as False Report Probability (FRP), and the *long-run probability* of H_1_ being true given a statistically significant result as True Report Probability (TRP). FRP and TRP are represented in **Row 5 of Table 1** and it is important to see that they refer to completely *different conditional probabilities than* the p value.

Simply put, the p value is pretty much the only thing that NHST computes but scientists usually would like to know the probability of their theory being true or false in light of their data (Jaynes, 2003; Pollard and Richardson, 1987; Goodman 1993). That is, researchers are interested in the post-experimental probability of H_0_ and H_1_. Most probably, for the reason that researchers do not get what they really want to see and the only parameter NHST computes is the p value it is well-documented (Oakes, 1986; Gliner et al. 2002; Wilkerson and Olson, 2010; Hoekstra et al. 2014; Castro-Sotos 2007; 2009) that many, if not most researchers confuse FRP with the p value or α and they also confuse the complement of p value (1-p) or α (1 -α) with TRP (Pollard and Richardson, 1987; Cohen 1994). These confusions are of major portend because the difference between these completely different parameters is not minor, they can differ by orders of magnitude, the long-run FRP being much larger than the p value under realistic conditions (Sellke et al. 2001; Ioannidis 2005). The complete misunderstanding of the probability of producing false positive findings is most probably a key factor behind vastly inflated confidence in research findings and we suggest that this inflated confidence is an important contributor to the current replication crisis in biomedical science and psychology.

### 3.1 Serious underestimation of the proportion of false positive findings in NHST

Ioannidis (2005) has shown that most published research findings relying on NHST are likely to be false. The modelling supporting this claim refers to the long-run FRP and TRP which we can compute by applying Bayes’ theorem (see computational details and illustrations in **Appendix 3**). The calculations must consider α, the power (1-β) of the statistical test used, the pre-study probabilities of H_0_ and H_1_, and it is also insightful to consider bias (Berger 1985; Berger and Sellke, 1987; Berger and Delampady, 1987; Sellke, Bayari and Berger, 2001; Pollard and Richardson, 1987; Lindley 1993; Sterne and Smith, 2001; Ioannidis 2005).

While NHST neglects the pre-study odds of H_0_ and H_1_, these are crucial to take into account when calculating FRP and TRP. For example, let's assume that we run 200 experiments and in 100 studies our experimental ideas are wrong (that is, we test true H_0_ situations) while in 100 studies our ideas are correct (that is, we test true H_1_ situations). Let’s also assume that the power (1-β) of our statistical test is 0.6 and α = 0.05. In this case in 100 studies (true H0) we will have 5% of results significant by chance alone and in the other 100 studies (true H_1_) 60% of studies will come up significant. FRP is the ratio of false positive studies to all studies which come up significant:

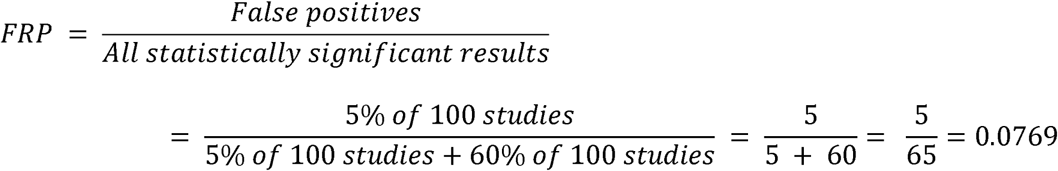

That is, we will have 5 false positives out of a total of 65 statistically significant outcomes which means that the proportion of false positive studies amongst all statistically significant results is 7.69%, higher than the usually assumed 5%. However, this example still assumes that we get every second hypothesis right. If we are not as lucky and only get every sixth hypothesis right then if we run 600 studies, 500 of them will have true H0 true situations and 100 of them will have true H_1_ situations. Hence, the computation will look like:

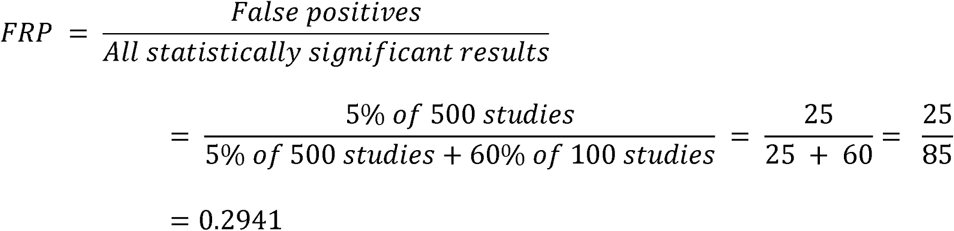

Hence, nearly 1/3 of all statistically significant findings will be false positives irrespective of the p value. Of course, estimating pre-study odds is difficult, primarily due to the lack of publishing negative findings and to the lack of proper documentation of experimenter intentions before an experiment is run: We do not know what percent of the published statistically significant findings are lucky false positives explained post-hoc when in fact researchers could not detect the originally hypothesized effect. However, it is reasonable to assume that only the most risk avoidant studies have lower H_0_:H_1_ odds than 1, relatively conservative studies have low to moderate H_0_:H_1_ odds (1-10) while H_0_:H_1_ odds can be much higher in explorative research (50-100 or even higher) (Ioannidis, 2005).

Bias is another important determinant of FRP and TRP (Ioannidis 2005). Whenever H_0_ is not rejected findings have far more difficulty to be published and the researcher may feel that she wasted her efforts. Further, positive findings are more likely to get cited than negative findings (Kivimäki et al. 2014; Jannot et al. 2013; Kjaergard and Gluud, 2002). Consequently, researchers may often be highly biased to reject H_0_ and publish positive findings. Researcher bias affects FRP even if our NHST decision criteria, α and β, are formally unchanged.

Ioannidis (2005) introduced the *u* bias parameter. The impact of u is that after some data tweaking and selective reporting (see **4.6**) u fraction of otherwise non-significant true H_0_ results will be reported as significant and u faction of otherwise non-significant true H_1_ results will be reported as significant. If u increases, FRP increases and TRP decreases. For example, if α = 0.05, power = 0.6 and H0:H1 odds = 1 then a 10% bias (u = 0.1) will raise FRP to 18.47%. A 20% bias will raise FRP to 26.09%. If H_0_:H_1_ odds = 6 then FRP will be 67.92%. Looking at these numbers the replication crisis does not seem surprising: using NHST very high FRP can be expected even with modestly high H_0_:H_1_ odds and moderate bias (Etz and Vanderckhove, 2016). Hence, under realistic conditions FRP not only *extremely rarely* equals α or the p value (and TRP extremely rarely equals 1-α and/or 1-p value) but also, FRP is *much* larger than the generally assumed 5% and TRP is much lower than the generally assumed 95%. Overall, α or the p value practically says nothing about the likelihood of our research findings being true or false.

### 3.2 The neglect of power reinterpreted

In contrast to the importance of power in determining FRP and TRP, NHST studies tend to ignore power and β and emphasize α and low p values. Often, finding a statistically significant effect erroneously seems to override the importance of power. However, statistical significance does not protect us from false positives. FRP can only be minimized by keeping H_0_:H_1_ odds and bias low and power high (Button et al. 2013; Pollard and Richardson, 1987; Bayarri et al. 2016). Hence, power is not only important so that we increase our chances to detect true effects but it is also crucial in keeping FRP low. While power in principle can be adjusted easily by increasing sample size, power in many/most fields of biomedical science and psychology has been notoriously low and the situation has not improved much during the past 50 years (Button et al. 2013; Cohen 1962; Sedlmeier and Gigerenzer, 1989; Rossi, 1990; Hallahan and Rosenthal, 1996). Clearly, besides making sure that research funding is not wasted, minimizing FRP also provides very strong rationale for increasing the typically used sample sizes in studies.

## 4. NHST logic is incomplete

### 4.1 NHST misleads because it neglects pre-data probabilities

Besides creating conceptual confusion and generating misleading inferences especially in the setting of weak power, NHST has further serious problems. NHST logic is based on the so-called *modus tollens* (denying the consequent) argumentation: It sets up a H_0_ model and assumes that if the data fits this model than the test statistic associated with the data should not take more extreme values than a certain threshold (Meehl, 1967; Pollard and Richardson, 1987). If the test statistic contradicts this expectation then NHST assumes that H_0_ can be rejected and consequently its complement, H_1_ can be accepted. While this logic may be able to minimize Type I error in well-powered high-quality well-controlled tests (**2.2**), it is inadequate if we use it to decide about the truth of H_1_ in a single experiment, because there is always space for Type I and Type II error (Falk and Greenbaum, 1995). So, our conclusion is never certain and the only way to see how much error we have is to calculate the long-run FRP and TRP using appropriate α and power levels and prior H_0_:H_1_ odds. The outcome of the calculation can easily conflict with NHST decisions (see **Appendix 4**).

### 4.2 NHST neglects predictions under H1 facilitating sloppy research

NHST does not require us to specify exactly what data H_1_ would predict. Whereas the Neyman-Pearson approach requires researchers to specify an effect size associated with H_1_ and compute power (1-β), in practice this is easy to *neglect* because the NHST machinery only computes the p value conditioned on H_0_ and it is able to provide this result even if H_1_ is not specified at all. A widespread *misconception* flowing from the fuzzy attitude of NHST to H_1_ is that rejecting H_0_ allows for accepting a *specific* H_1_ (Nickerson 2000). This is what most practicing researchers do in practice when they reject H_0_ and argue for their specific H_1_ in turn.

However, NHST only computes probabilities conditional on H_0_ and it does not allow for the acceptance of either H_0_, a specific H_1_ or a generic H_1_. Rather, it only allows for the rejection of H_0_. Hence, if we reject H_0_ we will have no idea about how well our data fits a specific H_1_. This cavalier attitude to H_1_ can easily lead us astray even when contrasting H_0_ just with a single alternative hypothesis as illustrated by the invalid inference based on NHST logic in **Table 2** (Pollard and Richardson, 1987).

Our model says that if H_0_ is true, it is a *very rare* event that Harold is a member of congress. This rare event then happens which is equivalent to finding a small p value. Hence, we conclude that H_0_ can be rejected and H_1_ is accepted. However, if we carefully explicate all probabilities it is easy to see that we are being mislead by NHST logic. First, because we have absolutely no idea about Harold’s nationality we can set pre-data probabilities of both H_1_ and H_0_ to 1/2, which means that H_0_:H_1_ odds are uninformative, 1:1. Then we can explicate the important conditional probabilities of the data (Harold *is* a member of congress) given the possible hypotheses. We can assign arbitrary but plausible probabilities:

> pr(data|H_0_) = pr(Harold is member of congress | American) = 10^−7^
>
> pr(data|H_1_) = pr(Harold is member of congress | not American) = 0

That is, while the data is indeed rare under H_0_, its probability is actually zero under H_1_ (in other words, the data is very unlikely under both the null and the alternative models). So, even if p ≈ 0.0000001, it does not make sense to reject H_0_ and accept H_1_ because this data just cannot happen if H_1_ is true. If we only have these two hypotheses to choose from then it only makes sense to accept H_0_ because the data is still possible under H_0_ (Jaynes, 2003). In fact, using Bayes' theorem we can formally show that the probability of H_0_ is actually 1 (**Appendix 5**).

In most real world problems multiple alternative hypotheses compete to explain the data. However, by using NHST we can only reject H0 and argue for *some* H_1_ without any formal justification of why we prefer a particular hypothesis whereas it can be argued that it only makes sense to reject any hypothesis if another one better fits the data (Jaynes, 2003). We only have qualitative arguments to accept a specific H_1_ and the exclusive focus on H_0_ makes unjustified inference too easy. For example, if we assume that H_0_ predicts normally distributed data with mean 0 and standard deviation 1 then we have endless options to pick H_1_ (Hubbard and Bayarri, 2003): Does H_1_ imply that the data have a mean other than zero, the standard deviation other than 1 and/or does it represent non-normally distributed data? NHST allows us to consider any of these options *implicitly* and then accept one of them post-hoc without any quantitative justification of why we chose that particular option. Further, merging all alternative hypotheses into a single H_1_ is not only too simplistic for most real world problems but it also poses an ‘inferential double standard’ (Rozeboom, 1960): The procedure pits the well-defined H_0_ against a potentially infinite number of alternatives.

Vague H_1_ definitions (the lack of quantitative predictions) enable researchers to avoid the falsification of their favourite hypotheses by intricately redefining them (especially in fields such as psychology and cognitive neuroscience where theoretical constructs are often vaguely defined) and ever providing any definitive assessment of the plausibility of a favourite hypothesis in light of credible alternatives (Meehl, 1967). This problem is reflected in papers aiming at the mere demonstration of often little motivated significant differences between conditions (Giere, 1972) and post-hoc explanations of likely unexpected but statistically significant findings. For example, neuroimaging studies often attempt to explain why an fMRI BOLD signal ‘deactivation’ happened instead of a potentially more reasonable looking ‘activation’ (or, vice versa). Most such findings may be the consequence of the data randomly deviating into the wrong direction relative to zero between-condition difference. Even multiple testing correction will not help such studies as they still rely on standard NHST just with adjusted α thresholds. Similarly, patient studies often try to explain an unexpected difference between patient and control groups (e.g. the patient group is ‘better’ on a measure) by some kind of ‘compensatory mechanism’. In such cases what happens is that *‘the burden of inference has been delegated to the statistical test’* indeed, and simply because p ≤ α odd looking observations and claims are to be trusted as scientific facts (Bakan, 1966, p423; Lykken 1968).

Finally, paradoxically, when we achieve our goal and successfully reject H_0_ we will actually be left in complete existential vacuum because during the rejection of H_0_ NHST ‘*saws off its own limb’*(Jaynes, 2003; p524): If we manage to reject H_0_ then it follows that pr(data or more extreme data|H_0_) is useless because H_0_ is not true. Thus, we are left with nothing to characterize the probability of our data in the real world; we will not know pr(data|H_1_) for example, because H1 is formally undefined and NHST never tells us anything about it. In light of these problems Jaynes (2003) suggested that the NHST framework addresses an ill-posed problem and provides invalid responses to questions of statistical inference.

### 4.4 The p value may exaggerate evidence against H0

The definition of the p value as pr(data or more extreme data | H_0_) is only justified informally by claiming that the p value is a measure of the ‘surprise’ felt when a rare event happens (Berger and Delampady, 1987). However, as we have seen our surprise at a rare event does not guarantee that H_0_ can be rejected: it may be surprising to find a member of Congress but this does not make him/her less likely to be American. Second, it can be shown formally that the definition of the p value does exaggerate the evidence against H_0_ by about one order of magnitude which greatly biases NHST procedures towards the rejection of H_0_ (see Berger and Sellke, 1987; Berger and Delamdapy, 1987; Goodman, 1993).

### 4.5 NHST is unsuitable for large datasets

In consequence of the recent ‘big data’ revolution access to large databases has increased dramatically potentially increasing power tremendously. However, NHST leads to worse inference with large databases than with smaller ones (Meehl, 1967; Khoury and Ioannidis, 2014). This is due to how NHST tests statistics are computed, the properties of real data and to the lack of specifying data predicted by H_1_ (Bruns and Ioannidis, 2016).

Most NHST studies rely on nil null hypothesis testing which means that H_0_ expects a true mean difference of exactly zero between conditions with some variation around this true zero mean. Further, NHST machinery guarantees that we can detect any tiny irrelevant effect sizes if sample size is large enough. This is because test statistics are typically computed as the ratio of the relevant between condition differences and associated variability of the data weighted by some function of the sample size (difference/variability × f(sample size)). The p value is smaller if the test statistic is larger. Thus, the larger is the difference between conditions and/or the smaller is variability and/or the larger is the sample size the larger is the test statistic and the smaller is the p value. Consequently, by increasing sample size enough it is guaranteed that H_0_ can be rejected even with miniature effect sizes (Ziliak et al. 2008).

Parameters of many real data sets are much more likely to differ than to be the same for reasons completely unrelated to our hypotheses (Edwards, 1972; Meehl, 1967; 1990). First, many psychological, social and biomedical phenomena are extremely complex reflecting the contribution of very large numbers of interacting (latent) factors, let it be at the level of society, personality or heavily networked brain function or other biological networks (Lykken 1968; Gelman, 2014). Hence, if we select any two variables related to these complex networks most probably there will be some kind of at least remote connection between them. This phenomenon is called ‘crud factor’ Meehl (1990) or ‘ambient correlational noise’ (Lykken, 1968) and it is unlikely to reflect a causal relationship. In fact some types of variables, such as intake of various nutrients and other environmental exposures are very frequently correlated among themselves and with various disease outcomes without this meaning that they have anything to do with causing disease outcomes (Patel and Ioannidis, 2014a,b). Second, unlike in physical sciences it is near impossible to control for the relationship of all irrelevant variables which are correlated with the variable(s) of interest (Rozeboom 1960; Lykken 1968).

Consequently, there can easily be a small effect linking two randomly picked variables even if their statistical connection merely communicates that they are part of a vast complex interconnected network of variables. Only a few of these tiny effects are likely to be causal and of any portend (Siontis and Ioannidis, 2011).

The above issues have been demonstrated empirically and by simulations. For example, Bakan (1966; see also Meehl, 1967; Nunally, 1960; Berkson, 1938) subdivided the data of 60,000 persons according to completely arbitrary criteria, like living east or west of the Mississippi river, living in the north or south of the USA, etc. and found all tests coming up statistically significant. Waller (2004) examined the personality questionnaire data of 81,000 individuals to see how many randomly chosen directional null hypotheses can be rejected. If sample size is large enough, 50% of directional hypothesis tests should be significant irrespective of the hypothesis. As expected, nearly half (46%) of Waller’s (2004) results were significant. Simulations suggest that in the presence of even tiny residual confounding (e.g. some omitted variable bias) or other bias, large observational studies of null effects will generate results that may be mistaken as revealing thousands of true relationships (Bruns and Ioannidis, 2016). Experimental studies may also suffer the same problem, if they have even minimal biases.

Due to the combination of the above properties of some psychological data sets and statistical machinery theory testing radically differs in sciences with exact and non-exact quantitative predictions (Meehl, 1967). In physical sciences increased measurement precision and increased amounts of data increase the difficulties a theory must pass before it is accepted. This is because theoretical predictions are well defined, numerically precise and it is also easier to control measurements (Lykken, 1968). That is, a theory may predict that a quantity should exactly be let’s say 5 and the experimental setup can assure that really only very few factors influence measurements - these factors can then be taken into account during analysis. Hence, increased measurement precision will make it easier to demonstrate a departure from numerically exact predictions. So, a ‘five sigma’ deviation rule may make good sense in physics where precise models are giving precise predictions about variables.

In sciences using NHST without clear numerical predictions the situation is the opposite of the above, because NHST does not demand the exact specification of H_1_, so theories typically only predict a fairly vague *‘difference’* between groups or experimental conditions rather than an exact numerical discrepancy between measures of groups or conditions. However, as noted, groups are actually likely to differ and if sample size increases and variability in data decreases it will become easier and easier to reject any kind of H0 when following the NHST approach. In fact, with precise enough measurements and large enough sample size H_0_ is guaranteed to be rejected on the long run even if the underlying processes generating the data in two experimental conditions are exactly the same. Hence, ultimately any H_1_ can be accepted, claiming support for any kind of theory. For example, in an amusing demonstration Carver (1993) used Analysis of Variance to re-analyze the data of Michelson and Morley (1887) who suggested that the speed of light is constant (H_0_) thereby providing the empirical basis for Einstein’s theory of relativity. Carver (1993) found that that the speed of light was actually not constant at p<0.001. The catch? The effect size as measured by Eta^2^ was 0.005. While some may feel that Einstein’s theory has now been falsified, perhaps it is also worth considering that here the statistically significant result is essentially insignificant.

A typical defence of NHST may be that we actually may not want to increase power endlessly, just as much as we still think that it allows us to detect reasonable effect sizes (Giere, 1972). A more reasoned approach may be to consider explicitly what the consequences (“costs”) are of a false-positive, true-positive, false-negative, and true-negative result. Explicit modelling can suggest that the optimal combination of Type 1 error and power may need to be different depending on what these assumed costs are (Ioannidis, Hozo and Djulbegovic, 2014). Different fields may need to operate at different optimal ratios of false-positives to false-negatives (Ioannidis, Tarone and McLaughlin, 2011).

### 4.6 NHST may foster selective reporting

Because NHST never evaluates H_1_ formally and it is fairly biased towards the rejection of H_0_, reporting bias against H_0_ can easily infiltrate the literature even if formal NHST parameters are fixed (see about the ‘u’ bias parameter in 3.1;). Overall, a long series of exploratory tools and questionable research practices are utilized in search for statistical significance (Johns 2012, Ioannidis and Trikalinos, 2007). Researchers can influence their data during undocumented analysis and pre-processing steps and by the mere choice of structuring the data (constituting *researcher degrees of freedom;* Simmons et al. 2011). This is particularly a problem in neuroimaging where the complexity and idiosyncrasy of analyses is such that it is usually impossible to replicate exactly what happened and why during data analysis (Carp 2012; Vul et al. 2009; Kriegeskorte et al. 2009). Another term that has been used to describe the impact of diverse analytical choices is “vibration of effects” (Ioannidis, 2008). Different analytical options, e.g. choice of adjusting covariates in a regression model can result in a cloud of results, instead of a single result, and this may entice investigators to select a specific result that is formally significant, while most analytical options would give non-significant results or even results with effects in the opposite direction (‘Janus effect’; Patel, Burford and Ioannidis, 2015). Another common mechanism that may generate biased results with NHST is when investigators continue data collection and re-analyse the accumulated data sequentially without accounting for the penalty induced by this repeated testing (DeMets and Lan, 1994; Goodman 1999). The unplanned testing is usually undocumented and researchers may not even be conscious that it exposes them to Type I error accumulation. Bias may be the key explanation why in most biomedical and social science disciplines, the vast majority of published papers with empirical data report statistically significant results (Fanneli 2010; Kavvoura et al. PLoS Med 2007; Chavalarias et al. 2016).

### 4.7 The rejection of H_0_ is guaranteed on the long-run

If H_0_ is false, with α = 0.05, 5% of our tests will be statistically significant on the long-run. The riskier experiments we run, the larger are H_0_:H_1_ odds and bias and the larger is the long-run FRP (3.3). Coupled with the fact that a large number of unplanned tests may be run in each study and that negative results and failed replications are often not published, this leads to *‘unchallenged fallacies’* clogging up the research literature (Ioannidis, 2012; p1; Bakan 1966; Sterling 1959; Sterling et al. 1995). Moreover, such published false positive true H0 studies will also inevitably overestimate the effect size of the non-existent effects or of existent, but unimportantly tiny, effects (Schmidt, 1992; 1996; Sterling et al. 1995; Ioannidis 2008). These effects may even be confirmed by meta-analyses, because meta-analyses typically are not able to incorporate unpublished negative results (Sterling et al. 1995) and they cannot correct many of the biases that have infiltrated the primary studies.

Given that the predictions of H_1_ are rarely precise and that theoretical constructs in many scientific fields (including psychology and cognitive neuroscience) are often poorly defined, it is easy to claim support for a popular theory with many kinds of data falsifying H_0_ even if the constructs measured in many papers are just very weakly linked to the original paper, or not linked at all. Overall, the literature may soon give the impression of a steady stream of replications throughout many years. Even when “negative” results appear, citation bias may still continue to distort the literature and the prevailing theory may continue to be based on the “positive” results. Hence, citation bias may maintain prevailing theories even when they are clearly false and unfounded (Greenberg, 2009).

### 4.8 NHST does not facilitate systematic knowledge integration

Due to high FRP the contemporary research literature provides statistically significant ‘evidence’ for nearly everything (Schoenfeld and Ioannidis, 2012). Because NHST emphasizes all or none p value based decisions rather than the magnitude of effects, often only p values are reported for critical tests, effect size reports are often missing and interval estimates and confidence intervals are not reported. In an assessment of the entire biomedical literature in 1990-2015, 96% of the papers that used abstracts reported at least some p value below 0.05, while only 4% of a random sample of papers presented consistently effect sizes with confidence intervals (Chalalarias, Wallach and Ioannidis, 2016). However, oddly enough, the main NHST ‘measure of evidence’, the p value cannot be compared across studies. It is a frequent *misconception* that a lower p value always means stronger evidence irrespective of the sample size and effect size (Oakes, 1986; Schmidt, 1996; Nickerson, 2000). Besides the non-comparable p values, NHST does not offer any *formal* mechanism for systematic knowledge accumulation and integration (Schmidt, 1996) unlike Bayesian methods which can take such pre-study information into account. Hence, we end up with many fragmented studies which are most often unable to say anything formal about their favourite H_1_s (accepted in a qualitative manner). Methods do exist for the meta-analysis of p values (see e.g. Cooper, Hedges and Valentine, 2009) and these are still used in some fields. However, practically such meta-analyses still say nothing about the magnitude of the effect size of the phenomenon being addressed. These methods are potentially acceptable when the question is whether there is any non-null signal among multiple studies that have been performed, e.g. in some types of genetic associations where it is taken for granted that the effect sizes are likely to be small anyhow (Evangelou and Ioannidis, 2013).

## 5. The state of the art must change

### 5.1 NHST is unsuitable as the cornerstone of scientific inquiry in most fields

In summary, NHST provides *the illusion of certainty* through supposedly ‘objective’ binary accept/reject decisions (Cohen, 1997; Ioannidis 2012) based on practically meaningless p values (Bakan 1966). However, researchers usually never give any formal assessment of how well their theory (a specific H_1_) fits the facts and, instead of gradual model building (Gigerenzer 1988) and comparing the plausibility of theories, they can get away with destroying a strawman: they disprove an H_0_ (which happens inevitably sooner or later) with a machinery biased to disproving it without ever going into much detail about the *exact* behaviour of variables under *exactly* specified hypotheses (Kranz 1999; Jaynes 2003). NHST also does not allow for systematic knowledge accumulation. In addition, both because of its shortcomings and because it is subject to major misunderstandings it facilitates the production of non-replicable false positive reports. Such reports ultimately erode scientific credibility and result in wasting perhaps most of the research funding in some areas (Nosek 2015; Ioannidis 2005; Macleod, Michie, Roberts, Dirnagl, Chalmers, Ioannidis, Al-Shahi Salman, Chan and Glasziou, 2014).

NHST seems to survive for various reasons. First, it allows for the easy production of a large number of publishable papers (irrespective of their truth value) providing a response to publication pressure. Second, NHST seems deceptively simple: because the burden of inference (Bakan, 1966) has been delegated to the significance test all too often researchers’ statistical world view is narrowed to checking an inequality: is p ≤ 0.05 (Cohen, 1994)? After passing this test, an observation can become a ‘scientific fact’ contradicting the random nature of statistical inference (Gelman, 2014). Third, in biomedical and social science NHST is often falsely perceived as the *single* objective approach to scientific inference (Gigerenzer et al. 1989) and alternatives are simply not taught and/or understood.

We have now decades of negative experience with NHST which gradually achieved dominance in biomedical and social science since the 1930s (Gigerenzer et al. 1989). Critique of NHST started not much later (Jeffreys, 1939) and has been forcefully present since then (Rozeboom, 1960; Nunnaly, 1960; Eysenck 1960; Clark 1963; Jeffreys, 1961; Bakan 1966; Mehl 1967; Lykken 1968) and continues to-date (Wasserstein and Lazar, 2016). The problems are numerous, and as Edwards (1972, p179) concluded 44 years ago: *‘any method which invites the contemplation of a null hypothesis is open to grave misuse, or even abuse’.* Time has proven this statement and that problems are unlikely to go away. We suggest that that it is *really* time for change now.

### 5.2 If theory is weak we need to focus on estimating effect sizes and their uncertainty

In basic biomedical and psychology research we often cannot provide very well worked out hypotheses and even a simple directional hypothesis may seem particularly enlightening. Such rudimentary state of knowledge can be respected. However, in such pre-hypothesis stage substantively blind all or nothing accept/reject decisions may be unhelpful and may maintain our ignorance rather than facilitate organizing new information into proper scientific models. It is much more meaningful to focus on assessing the magnitude of effects along with estimates of uncertainty, let these be error terms or Bayesian credible intervals (Luce, 1988; Edwards, 1972; Jaynes 2003; Schmidt, 1996; Gelman 2013; Lakens & Evers, 2014; see Morey et al. 2016 for why classical confidence intervals are not appropriate uncertainty measures). These provide more direct information on the actual ‘empirical’d behaviour of our variables. Gaining enough experience with interval estimates and assuring their robustness by building replication into design (Nosek et al. 2013) may then allow us to describe the behaviour of variables by more and more precise scientific models which may provide more clear predictions (Jaynes 2003; Schmidt, 1996; Gelman 2013).

The above problem does not only concern perceived ‘soft areas’ of science where measurement, predictions, control and quantification are thought to be less rigorous than in ‘hard’ areas (Meehl, 1978). In many fields, for example, in cognitive neuroscience, the measurement methods may be ‘hard’ but theoretical predictions and analysis often may be just as ‘soft’ as in any area of ‘soft’ psychology: Using a state of the art fMRI scanner for data collection and extremely complicated and often not clearly understood black box software for data analysis will not make a badly defined theory well defined. In fact, in such *pretend-hard* areas the situation may even be worse because technology allows us to measure a huge number of variables and run an immense number of tests (many of them undocumented and hence, uncorrected for multiple testing) and analyze the data by highly complex obscure black box processes and non-replicable idiosyncratic approaches. All these problems will only boost the number of false positive unreplicable findings (Carp, 2012; Vul et al. 2009; Kriegeskorte et al. 2009).

### 5.3 Improved reporting and alternative statistical inference methods are needed

While we need substantial change, criticism of NHST should not *‘lapse into methodological anarchy out of despair or confusion’* (Giere, 1972, p. 171). For example, recently, the Journal of Basic and Applied Social Psychology banned NHST from their articles (Trafimov and Marks, 2015; see Hunter, 1997). The decision prompted critical responses from several high profile statisticians who objected to the approach of the journal editors and their negative view on the controversies of statistical inference in general (https://www.statslife.org.uk/news/2116-academic-journal-bans-p-value-significance-test), which sharply contrasts with the view of the renowned physicist ET Jaynes (2003; pp. xxii; see starting quotes) who plausibly defined probability theory as the *‘logic of science’.* Let‘s make it clear that we are not arguing against statistical inference in general and we do not want to ban NHST. Quantitative and well justified statistical inference should be at the *core* of the scientific enterprise. We argue against the *default* and mindless application of NHST.

It may be reasonable to use NHST in some cases. One such case is when very precise quantitative theoretical predictions can be tested, hence, both power and effect size can be estimated well as intended by Neyman and Pearson (1933). Further, when theoretical predictions are not precise, we powered NHST tests may be used as an initial heuristic look at the data as Fisher (1925) intended. In this second case NHST tests must be followed up by more robust procedures to estimate effect sizes and interval estimates (e.g. by the now widely available bootstrap and permutation procedures) and (if there are clear hypotheses) more robust likelihood estimation or Bayesian techniques to test hypotheses. As Fisher was well conscious, NHST procedures can only suggest that something is really important if they *‘rarely fail’* to give us statistically significant results (Gigerenzer et al. 1989, p96; Goodman, 2008). Hence, strong claims require the replication of NHST tests optimally within the initial study. These replications must be well powered to keep FRP low. In all cases when NHST is used its use must be justified clearly rather than used as an automatic default and single cornerstone procedure. As discussed, NHST can only reject H_0_ and can accept neither a generic or specific H_1_. So, on its own NHST cannot provide evidence ‘for’ something even if findings are replicated.

Statistical reporting must improve substantially. Optimally, raw data should be published because data parameters of interest depend on the choice of models and analyses. Regarding the actually chosen analyses, the distribution (very rarely plotted at the moment) and important parameters of the data should be communicated (e.g. means and standard deviations in the original units of measurement as well as confidence and/or credible intervals). Regarding NHST procedures, researchers should report power calculations for each test including those with non-significant results and the number of cases should clearly be reported for each test. In the age of internet all important results can be communicated as online supplementary material, so there is not much excuse for not doing this. Improved access to raw data, algorithms and code would also be helpful and efficient ways to promote such reproducible research practices need to be found (Doshi, Goodman, Ioannidis, 2013; Diggle and Zeger, 2010; Keiding 2010; Laine, Goodman and Griswold, 2007; Peng, 2009; Peng, 2011). Incentives such as a badge system may help promote availability of more raw data (Nosek et al. 2015). Diverse stakeholders (journals, funders, institutions, and more) may contribute to align incentives with better research practices (Ioannidis 2014).

Hypotheses could be tested by either likelihood ratio testing, and/or Bayesian methods which usually view probability as characterizing the state of our beliefs about the world (Jaynes, 2003; Pearl 1998; MacKay, 2003; Gelman et al. 2014; Sivia and Skilling, 2006). The above alternative approaches require model specifications about alternative hypotheses, they can give probability statements about H_0_ and alternative hypotheses, they allow for clear model comparison, are insensitive to data collection procedures and do not suffer from problems with large samples. In addition, Bayesian methods can also factor in pre-study (prior) information into model evaluations which may be important for integrating current and previous research findings. Hence, the above alternative approaches seem more suitable for the purpose of scientific inquiry than NHST and ample literature is available on both. The problem is that usually none of these alternative approaches are taught properly in statistics courses for students in psychology, neuroscience, biomedical science and social science. For example, across 1000 abstracts randomly selected from the biomedical literature of 1990-2015, none reported results in a Bayesian framework (Chavalarias, et al. 2016).

### 5.4 Better training and better use of statistical methods: from believers to thinkers

We argue that the practice of relying on (good-willed) editorial dictates rather than informed statistical thinking is a symptom of a core problem: the statistical subject knowledge of many researchers in biomedical and social science has been shown to be poor (Oakes, 1986; Gliner et al. 2002; Wilkerson and Olson, 2010; Hoekstra et al. 2014; Castro-Sotos 2007; 2009). NHST perfectly fits with poor understanding because of the perceived simplicity of interpreting its outcome: is p ≤ 0.05 (Cohen, 1994)?

We suggest that the weak statistical understanding is probably due to inadequate ‘statistics lite’ education based on supposedly ‘user friendly’ dumbed down statistics cookbooks which may do more harm than good. This approach does not build up appropriate mathematical fundamentals and does not provide scientifically rigorous introduction into statistics. Hence, students’ knowledge may remain imprecise, patchy and prone to serious misunderstandings. What this approach achieves, however, is providing students with false confidence of being able to use inferential tools whereas *they usually only interpret the p value provided by black box statistical software.* While this educational problem remains unaddressed, poor statistical practices will prevail regardless of what procedures and measures may be favoured and/or banned by editorials.

Understanding probability is difficult. Common sense is notoriously weak in understanding phenomena based on probabilities (Gigerenzer et al. 2005). We cannot assume that without proper training biomedical and social science graduates would get miraculously enlightened about probability. Some of the best symbolic thinking minds of humanity devoted hundreds of years to the proper understanding of probability and statisticians still do not agree on how best to draw statistical inference (Stigler 1986; Gigerenzer et al. 1989), e.g. the recent ASA statement on p values (Wasserstein and Lazar, 2016) was accompanied by 21 editorials from the statisticians and methodologists who participated in crafting it and who disagreed in different aspects among themselves.

One approach would be to phase out the ‘statistics lite education approach for all research stream students and teach statistics rigorously, based on two years of calculus. Besides NHST, Bayesian and likelihood based approaches should also be taught, with explanation of the strengths and weaknesses of each inferential method. An alternative and/or complementary approach would be to enhance the training of professional applied statisticians and to ensure that all research involves knowledgeable statisticians or equivalent methodologists. At a minimum, all scientists should be well trained in understanding evidence and statistics and being in a position to recognize that they may need help from a methodologist expert (Marusic A, Marusic, 2003; Moharar, Rahimi and Najafi, 2009; Vujaklija, Hren, Sambunjak, Vodopivec, Ivanis, Marusic and Marusic, 2010).

All too often statistical understanding is perceived as something external to the subject matter of substantive research. However, it is important to see that statistical understanding influences most decisions about substantive questions, because it underlies the *thinking* of researchers even if this remains *implicit.* While common sense ‘statistics’ may be able to cope with simple situations, common sense is not enough to decipher scientific puzzles involving dozens, hundreds, or even thousands of interrelated variables. In such cases well justified applications of probability theory are necessary (Jaynes, 2003). Hence, instead of delegating their judgment to ‘automatized’ but ultimately spurious decision mechanisms, researchers should have confidence in their own *informed judgment* when they make an inference.

### 5.4 There is no automatic inference: New-old dangers ahead?

Perhaps the most worrisome false belief about statistics is the belief in automatic statistical inference (Bakan 1966), the illusion that plugging in some numbers into some black box algorithm will give a number (perhaps the p value or some other metric) that conclusively proves or disproves hypotheses (Bakan 1966). There is no reason to assume that any kind of ‘new statistics’ (Cummings, 2008) will not suffer the fate of NHST if statistical understanding is inadequate. For example, it has been shown that confidence intervals are misinterpreted just as badly as p values by undergraduates, graduates and researchers alike and self-declared statistical experience even slightly positively correlates with the number of errors (Hoekstra et al. 2014). Similarly, the proper use of Bayesian methods may require use of advanced simulation methods and a clear understanding and justification of probability distribution models. In contrast to this, it is frequent to see a kind of ‘automatic’ determination of Bayes factors or posterior estimates, again, provided by black box statistical packages which again, promise to take the load of thinking off the shoulders of researchers.

There is no reason to assume that understanding 21^st^ and 22^nd^ century science will require less mathematical and statistical understanding than before. If statistical understanding does not improve it will not matter whether editorials enforce bootstrapping, likelihood estimation or Bayesian approaches, they will all remain mystical to the untrained mind and open to abuse such as the NHST of the 20^th^ century.

## Acknowledgments.

DS is supported by the James S McDonnell Foundation.

